# Whole human genome 5’-mC methylation analysis using long read nanopore sequencing

**DOI:** 10.1101/2021.05.20.444035

**Authors:** Catarina Silva, Miguel Machado, José Ferrão, Sebastião Rodrigues, Luís Vieira

**Affiliations:** Unidade de Tecnologia e Inovação, Departamento de Genética Humana, Instituto Nacional de Saúde Doutor Ricardo Jorge, Av. Padre Cruz, 1649-016 Lisboa, Portugal; Centro de Toxicogenómica e Saúde Humana (ToxOmics), Genética, Oncologia e Toxicologia Humana da Nova Medical School|Faculdade de Ciências Médicas, Universidade Nova de Lisboa, Campo dos Mártires da Pátria 130, 1169-056 Lisboa, Portugal

## Abstract

DNA methylation is a type of epigenetic modification that affects gene expression regulation and is associated with several human diseases. Microarray and short read sequencing technologies are often used to study 5’-methylcytosine (5’-mC) modification of CpG dinucleotides in the human genome. Although both technologies produce trustable results, the evaluation of the methylation status of CpG sites suffers from the potential side effects of DNA modification by bisulfite and the ambiguity of mapping short reads in repetitive and highly homologous genomic regions, respectively. Nanopore sequencing is an attractive alternative for the study of 5’-mC since the long reads produced by this technology allow to resolve those genomic regions more easily. Moreover, it allows direct sequencing of native DNA molecules using a fast library preparation procedure. In this work we show that 10X coverage depth nanopore sequencing, using DNA from a human cell line, produces 5’-mC methylation frequencies consistent with those obtained by methylation microarray and digital restriction enzyme analysis of methylation. In particular, the correlation of methylation values ranged from 0.73 to 0.90 using an average genome sequencing coverage depth <2X or a minimum read support of 17X for each CpG site, respectively. We also showed that a minimum of 5 reads per CpG yields strong correlations (>0.89) between sequencing runs and an almost uniform variation in methylation frequencies of CpGs across the entire value range. Furthermore, nanopore sequencing was able to correctly display methylation frequency patterns according to genomic annotations, including a majority of unmethylated and methylated sites in the CpG islands and inter-CpG island regions, respectively. These results demonstrate that low coverage depth nanopore sequencing is a fast, reliable and unbiased approach to the study of 5’-mC in the human genome.

## Introduction

The transcriptional dynamics of a cell’s genome can be influenced by epigenetic mechanisms that do not involve modifying the DNA sequence. The set of epigenetic changes in a genome produces the epigenome, which differs from individual to individual. The epigenome includes different types of chemical modifications of DNA, such as methylation, alterations of histone proteins (e.g., methylation, acetilation), the expression of non-coding RNAs and higher-order changes in the chromatin structure^1^. One of the best studied epigenetic mechanisms is DNA methylation, a process that occurs post-replication in embryonic, germinal or somatic cells, and that spreads through mitosis. In mammals, the methylation pattern is established *de novo* at the beginning of development and in gametogenesis by DNA methyltransferases (DNMTs) DNMT3A and DNMT3B^2^. These enzymes transfer a methyl group from 5-adenyl methionine to the fifth carbon atom of the 5’-cytosine residue, forming 5’-methylcytosine (5’-mC). This epigenetic modification is highly conserved in eukaryotic organisms^3^. DNA methylation is a cyclic and dynamic enzymatic process that involves methylation of cytosines by DNMTs, oxidation of methyl groups by dioxygenase enzymes, and restoration of cytosines in their original form by active or passive demethylation mechanisms^3^.

DNA methylation frequently affects cytosines located in cytosine-phosphate-guanine (CpG) dinucleotides, and are rare in other sequence contexts in mature mammalian cells^3^. However, methylation activity is dependent on the sequence context in which the DNA modification occurs. If methylation occurs within the promoter regions of genes, it typically has a repressive effect on gene expression, while methylation at the level of gene bodies has a transcriptional activation role^4^. In regulatory regions, DNA methylation can prevent the binding of transcription factors, thereby repressing gene expression. Furthermore, methylation of cytosine residues can lead to recruitment of methyl-binding proteins, which in turn attract members of the chromatin remodeling complex such as histone deacetylases, leading to a change in the chromatin conformation at the local level that can have an effect on the activation or repression of gene expression depending on the cellular context^5^. CpG islands are often located on the promoters of genes and are characterized by a length of at least 500 bp with >55% GC content. CpG shores are genomic regions that extend at a maximum distance of 2 kilobases (kb) from the CpG islands and are characterized by tissue-specific differential methylation patterns. CpG shelves correspond to the genomic regions that are located between 2 and 4 kb distant from the CpG islands. The remaining genomic regions constitute the open sea regions and represent the vast majority of the human genome sequence.

The alteration of the DNA methylation pattern is associated with several human diseases. Some genetic diseases are related to genomic imprinting, a constitutional epigenetic mechanism that involves the methylation of DNA and histones, through which some genes are expressed only from the chromosome inherited from the paternal or maternal side^6^. Epigenetic mechanisms are also associated with the development of some common diseases such as autoimmune, cardiovascular, metabolic and psychiatric diseases^7–10^, in which genetic variants may not explain the totality of observed phenotypes and in which epigenetic therapies (e.g., DNA methylation targeting by CRISPR/Cas9 tools) may be beneficial. In myelodysplastic syndromes and acute myeloid leukemia, chemical demethylating agents (e.g., 5-azacytidine) are used in the treatment of patients, whose function is to suppress epigenetic marks and thus reactivate the normal expression of genes silenced during the tumorigenic process^11^. A relationship between environmental exposure (e.g., tobacco smoke) and methylation has already been shown for several compounds, leading to a change in the overall methylation profile (e.g., hypomethylation) or in the methylation of specific genes^12^. These data demonstrate the need for further studies on the role of human genome methylation in a disease context.

The study of 5’-mC profile was carried out, over many years, through chemical treatment of DNA with bisulfite, followed by amplification with strand-specific and bisulfite-specific primers, and sequencing by the dideoxynucleotide chain-termination method^13^. Bisulfite converts each unmethylated cytosine to uracil, while 5’-methylcytosine is resistant to conversion. During amplification, each uracil residue is amplified as thymine while the 5’-mC residues are amplified as cytosines, which allows the distinction of methylated cytosines from the unmethylated ones through sequencing of the amplified fragments. Although bisulfite DNA modification technique is widely used in different methylation study protocols^14^, chemical modification of DNA has several side effects on sequence structure and integrity that can potentially affect the results^15–17^. Due to the strong development that was witnessed in the microarray and next-generation sequencing technologies, and in the bioinformatics methods for high-throughput data analysis, the study of chemical changes at the DNA sequence level has moved from an approach focused on individual *loci* to a genome-wide scale in recent years^18–20^.

More recently, nanopore sequencing technology, which generates very long reads using native DNA molecules as input, has emerged as an alternative approach to the study of methylation. The electrical signals registered at the nanopore level are also sensitive to the presence of modifications in the DNA chain, and can be used to differentiate methylated cytosines from unmethylated cytosines using a hidden Markov model^21^. However, there are still very few studies in which nanopore sequencing was used to analyze the 5’-mC profile in the human genome^21–23^. In the present work, we describe the complete genome sequencing of a human cell line with nanopore technology, and present a benchmark comparison of 5’-mC methylation status compared to other traditional methods, in order to evaluate the performance of the technology in the quantification of methylation at CpG sites. This cell line had been previously studied for 5’-mC methylation using the Illumina Infinium HumanMethylation450 BeadChip microarray assay^24^, using bisulfite-treated DNA, and Digital Restriction Enzyme Analysis of Methylation (DREAM)^25^, a method based on short read sequencing of DNA fragments resulting from sequential digestion with a pair of neoschizomeric restriction enzymes that recognize the CCCGGG sequence. The 450k microarray allows the analysis of 485 512 CpG sites, which constitute ~1.7% of the existing CpGs in the human genomic sequence and that were selected based on the opinion of a panel of experts, with the main focus being the CpGs located in genes and in CpG islands^26^. The DREAM method allows to quantify 374 170 CpG sites, corresponding to ~1.3% of the CpGs present in the hg18 human genome reference sequence^25^, of which approximately 10% are located in CpG islands. Using low sequencing coverage depth (10X), we showed that the methylation frequencies of called CpG sites based on nanopore sequencing are strongly correlated with those obtained by microarray and DREAM. We conclude that this approach has a high potential for the study of 5’-mC changes in the human genome.

## Materials and methods

### Cell line

The human erythroleukemia (HEL) cell line, established from the peripheral blood of a patient with erythroleukemia developed after treatment for Hodgkin’s disease^27^, was purchased directly from DSMZ - German Collection of Microorganisms and Cell Cultures GmbH (Braunschweig, Germany). The cells were placed and maintained in a T25 standard culture flask, according to the supplier’s instructions and to standard protocols for culturing cell lines. After cell confluence was achieved, approximately one third of the culture was transferred into a freezer vial. The culture medium was removed by centrifugation and the cell pellet was ressuspended in a mixture of 90% fetal bovine serum and 10% DMSO. The cell suspension was primarily frozen at −80°C for 24 hours and then transferred to a liquid nitrogen container, where it remained until genomic DNA extraction.

### DNA extraction, quantification and quality analysis

The freezing vial was thawed completely in the hand and the entire volume of cell suspension (~1 ml) was transferred to a 2 ml-microtube. The microtube was centrifuged for 3 minutes at 1000 rpm to deposit the pellet and the supernatant was carefully aspirated with a micropipette and discarded. The cells were washed using 1 ml of Dulbecco’s Phosphate Buffered Saline 1X (Gibco) and centrifugation was repeated. The supernatant was discarded and the cell pellet was carefully ressuspended in 700 μl of sodium chloride-TRIS-EDTA buffer to which 35 μl of 10% sodium dodecyl sulfate (Sigma) and 7 μl of proteinase K at 10 mg/μl (AmpliChem) were added. The mixture was incubated at 55°C for 3 hours, with gently tapping the microtube from time to time. Then, 700 μl of aquaphenol water saturated (MP Biomedicals) were added and the mixture was manually stirred very slowly by inversion and centrifuged for 5 minutes at 13000 rpm. The aqueous phase was aspirated slowly using a micropipette with 200 μl-tip. The recovered volume was divided by 2 tubes and the centrifugation was repeated with 700 μl of aquaphenol. The aqueous phase recovered from each tube was transferred to a new 2 ml-microtube. After the addition of 700 μl of chloroform p.a. (Merck), the microtube was slowly and manually inverted to mix. The mixture was then centrifuged 5 minutes at 13000 rpm, after which the aqueous phase was removed into a new 2 ml-microtube where 70 μl of 3M sodium acetate and 1.4 ml of ice-cold 100% ethanol were added. The mixture was then inverted several times very slowly, to avoid shearing, until the strands of genomic DNA were noticeable. After centrifugation at 4°C for 15 minutes at 13000 rpm, the supernatant was discarded by aspiration and 1 ml of ice-cold 70% ethanol was added to wash the DNA pellet. The mixture was gently stirred by inversion and centrifuged at 4°C for 15 minutes at 13000 rpm. The supernatant was discarded by inverting the microtube. The precipitated genomic DNA was dried at room temperature by placing the inverted microtube on absorbent paper for 15 minutes. After drying, the DNA was ressuspended in 150 μl of low Tris-EDTA buffer and placed in a dry bath at 55°C without shaking for 1 hour. Then, the microtube was transferred to 4°C, with stirring by inversion from time to time, and kept overnight at this temperature. The volume was then transferred to a 1.5-ml MAXYMum recovery microtube (Axygen). DNA was quantified by spectrophotometry and fluorimetry using NanoDrop One and Qubit 3.0 (ThermoFisher Scientific), respectively. The DNA sample was diluted to a final concentration of 70 ng/μl. A total of 140 ng of genomic DNA was electrophoresed on a 0.8% agarose gel stained with ethidium bromide, and compared with the lambda molecular weight marker DNA/Hind III (Invitrogen). Genomic DNA was placed at −20°C in 40 μl aliquots for long-term storage.

### DNA dialysis

An aliquot (40 μl) of genomic DNA was subjected to purification by dialysis, using a method previously described^28^ with minor adaptations, to try to increase the purity of the sample prior to sequencing. After thawing at room temperature, the aliquot was placed in a dry bath for 15 minutes at 55°C, with slight stirring every 5 minutes, to homogenize the solution. Dialysis was performed using a 25 mm sterile acrodisc^®^ syringe filters with supor^®^ membrane with 0.2 μm of pore size (PALL). The filter was suspended in 4 ml of sterile bidistilled water within the plastic wrapper, immersed up to the membrane level, with the part that serves as the fitting for the syringe oriented to the outside. After 5 minutes of moistening the membrane, the entire volume of the aliquot was pipetted into the syringe fitting part and the inlet of the filter was subsequently covered to prevent evaporation. After 1 hour of dialysis, and with the filter in place, 35 μl of dialyzed DNA solution (87.5% recovery) was aspirated into a microtube.

### Library preparation and sequencing

Before preparing the libraries, genomic DNA was placed for 15 minutes at 55°C in the dry bath without shaking. Four genomic libraries were prepared using the Rapid Sequencing Kit SQK-RAD004 (Oxford Nanopore Technologies, ONT). Two libraries were prepared with an initial DNA input of 400 ng, as indicated by the manufacturer, whereas the other two were prepared with an input of 280 ng and 1500 ng (7.5 μl of solution after vacuum drying). Sequencing was performed on the MinION device (ONT) using FLO-MIN106D flow cells. Data was transferred via USB connection to a portable computer with an Intel i7 processor and 16 GB of RAM. The MinKNOW software (MinION Release 19.06.8, ONT) was used to program and configure run parameters, and to acquire the data (FAST5 files) during sequencing.

### Sequencing data processing and analysis

The FAST5 files were transferred to a Linux server with 48 CPUs and 384 GB of RAM. The raw data was analyzed with NanoPlot (v1.30.1)^29^ to obtain various run statistics. Methylation analysis was performed using the Nanopype pipeline (v1.0.0)^30^. Briefly, basecalling was carried out with Guppy (ONT)^31^, read alignment was performed using ngmlr^32^ and minimap2^33^ against the hg38 reference human genome sequence (Genome Reference Consortium Human Build 38 patch 12 - GRCh38.p12, Ensembl Release 96) and the calling of 5’-mC was performed with nanopolish^21^. The Nanopype pipeline was applied to each run independently (using minimap2-base mappings only) and to the combined data of the 4 runs (using ngmlr and minimap2 mappings). A total of 6 ‘frequencies.tsv’ files containing methylation frequencies (corresponding to the ratio between the number of methylated sites and the number of called sites) for each mapped CpG dinucleotide were obtained. Coverage statistics were obtained with the bamqc tool of Qualimap2^34^ software (v2.2.1), after merging, sorting and indexing the BAM files with samtools^35^ (v1.9).

### Comparative analysis of methylation frequencies

The methylation frequencies obtained using nanopore sequencing were compared with previously available data of methylation microarray^24^ and DREAM^25^ for the HEL cell line. The microarray and DREAM data are available in the Gene Expression Omnibus series records GSE68379 and GSE39787 for the HEL cell line (samples GSM1669874 and GSM979022, respectively). The LiftOver tool^36^ was used to convert the genomic coordinates of the DREAM data from hg18 to hg19, and from hg19 to hg38, and from the microarray data from hg19 to hg38 genome builds. The genomic positions of CpGs in the microarray and DREAM data were corrected to n-1 and n-2, respectively, for matching with the positions obtained in the methylation frequency files generated by nanopolish, and then randomly confirmed between all files using the flanking sequences of the CpG sites. The methylation frequency in the DREAM dataset was calculated by dividing the number of methylated reads by the total number of reads for each CpG site. The BEDTools^37^ (v2.27.1) intersect function was used to intersect the coordinates of the CpG sites between the various data files. The resulting intersection files were analyzed with several R packages (V4.0.2). The ‘corrr’ (v0.4.3)^38^ and ‘corrgram’ (v1.13)^39^ packages were used to calculate the product-moment Pearson correlation between the methylation values obtained in nanopore sequencing, DREAM, and microarray. The data files obtained by nanopore sequencing were also filtered according to a minimum read support per CpG site, using the sed and awk commands in the Linux shell, and the resulting files were then intersected with the methylation microarray data file to assess the correlation of methylation values for common CpG sites. The genomic annotations of CpG sites (CpG islands, shores, shelves and open sea regions) were obtained for chromosome 1 using the ‘annotatr’^40^(version 1.14.0) R package.

## Results

### Sample and sequencing quality analysis

The genomic DNA of the HEL cell line showed a single band with a minimum molecular size of 23 kb and no evidence of smear. The 260nm/280nm ultra-violet absorption ratio was identical (1.95) in the sample without dialysis and in the dialyzed sample, while the 260nm/230nm ratios were, respectively, 2.06 and 2.14, indicating a potential benefit of dialysis to remove traces of phenol. Three nanopore sequencing runs were performed with DNA without dialysis (runs 1 to 3) and one with dialyzed DNA (run 4). The 4 runs had a total yield of 33.31 Gb (ranging between 5.66 and 12.59 Gb per run), encompassing a total of 4 249 514 reads. The longest read had a length of 292471 base pairs (bp). The read length N50 was 14.94 kb and the mean and median of the length of the reads were 7841 bp and 4366 bp, respectively (**Table I**). The mean read quality value (QV) was of 10.1 and the majority of the reads (67.4%) had a very high quality (QV>10) (**Figure 1**). The overall quality decreased slightly during the run, but most of the bases sequenced at the end of the runs still showed a high quality. The mean read quality was higher in run 4 (10.3) compared to any of the other runs (10), which may be attributed to the dialysis procedure. These results demonstrate that the quality of the DNA prepared by the phenol-chloroform method associated with the rapid method of library preparation, allows the generation of a large amount of high quality sequencing data.

**Table I.** Sequencing run statistics.

| Run number |  | 1 | 2 | 3 | 4 | Combined |
| --- | --- | --- | --- | --- | --- | --- |
| DNA input (ng) |  | 280 | 400 | 1500 | 400 |  |
| Starting pores available for sequencing |  | 1701 | 1462 | 1478 | 1692 |  |
| Run time (H) |  | 45 | 48 | 48 | 48 |  |
| Before base calling | Number of Reads (K) | 465 | 730 | 1440 | 1620 |  |
|  | Output (Gb) | 5.91 | 7.46 | 7.83 | 12.78 |  |
|  | N50 (Kb) | 23.67 | 21.33 | 10.02 | 13.73 |  |
|  | Longest Read (Kb) | 257 | 305 | 196 | NA |  |
| After base calling | Number of Reads | 465 108 | 729 775 | 1 435 541 | 1 619 090 | 4 249 514 |
|  | Output (Gb) | 5.66 | 7.24 | 7.82 | 12.59 | 33.31 |
|  | N50 (Kb) | 23.15 | 18.58 | 10.22 | 13.71 | 14.94 |
|  | Longest Read (pb) | 240905 | 292471 | 185813 | 186627 | 292471 |
|  | Mean read length (pb) | 12167 | 9928 | 5452 | 7778 | 7841 |
|  | Median read length (pb) | 6998 | 5639 | 3015 | 4836 | 4366 |
|  | Mean read quality | 10 | 10 | 10 | 10.3 | 10.1 |
|  | Median read quality | 10.8 | 10.7 | 10.7 | 10.9 | 10.8 |
|  | % QV>7 | 87.6 | 89 | 89.2 | 90.5 | 89.5 |
|  | % QV>10 | 66.7 | 65.9 | 64.8 | 70.5 | 67.4 |

NA, not available

**Figure 1.**
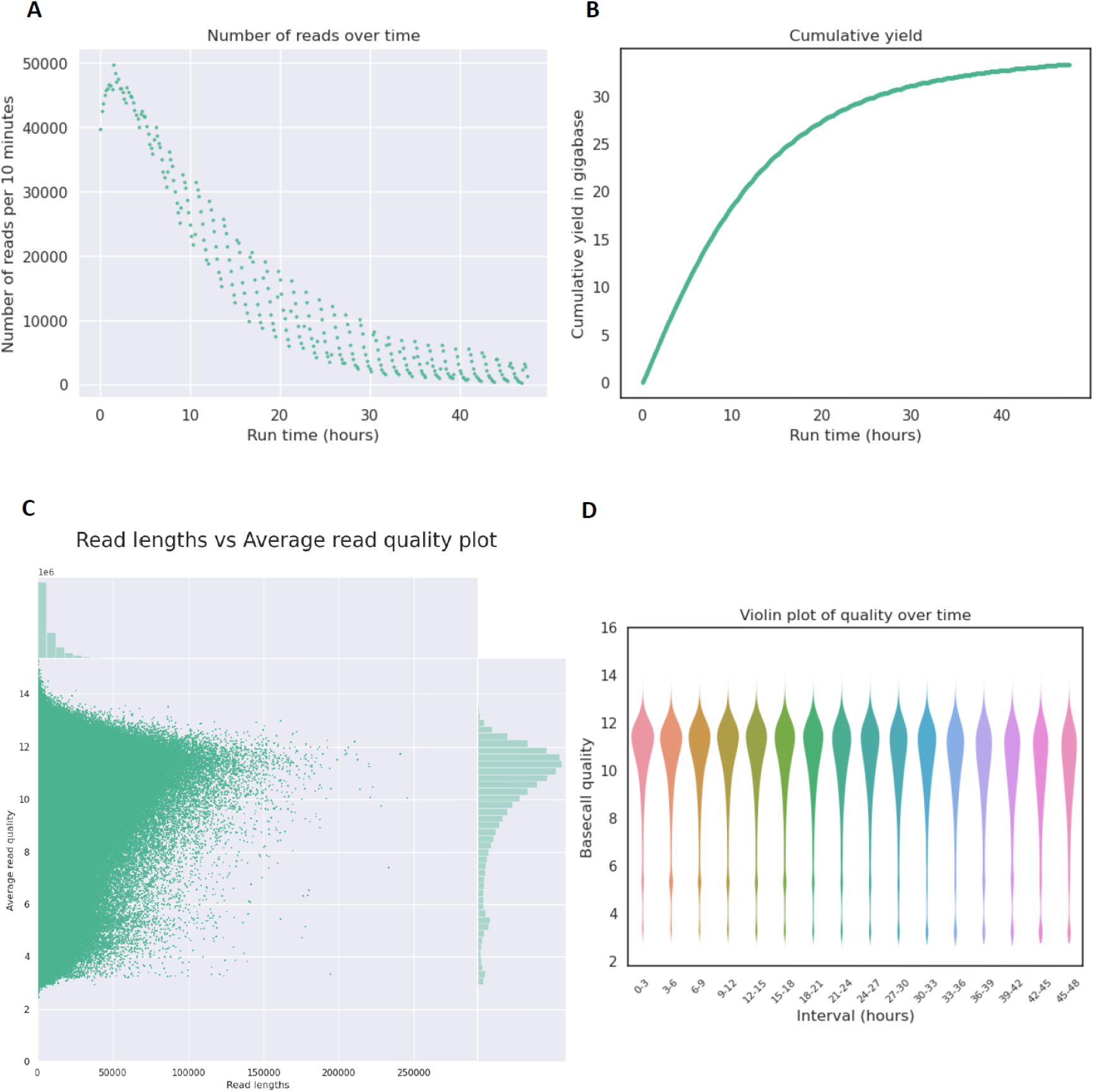
Quality analysis of nanopore sequencing reads generated using genomic DNA from the HEL cell line. The plots represent various quality metrics for the combined set of reads produced in 4 sequencing runs: **A**, number of reads over run time; **B**, cumulative yield (in gigabases) over run time; **C**, read length versus average read quality; **D**, basecall quality over run time. The majority of reads had an average read quality between 10 and 14. After 48 hours of sequencing run, there were still new reads being produced, the majority of these having a basecall quality over 10. Plots were generated using Nanoplot (v1.30.1).

### Genomic coverage and annotation of CpG sites

The reads of the HEL cell line mapped in the hg38 reference sequence of the human genome, using minimap2 and ngmlr, constituted 93.6% and 88.6% of the total reads, which resulted in an average coverage depth of 10.6X (standard deviation: 28.6X) and 10.0X (standard deviation: 20.9X), respectively. Methylation frequencies were obtained for a total of 29.093.016 and 28.114.726 CpG dinucleotides respectively, despite the reduced depth of coverage. These results indicate that it was possible to obtain information on the status of 5’-mC in practically all CpGs (~28M) present in the hg19 human genome reference sequence^41^. The genomic annotation of the CpG dinucleotides located in the chromosome 1 sequence, revealed the expected pattern according to the predicted length of the annotated regions, and characterized by a similar proportion of CpG dinucleotides located in the CpG islands, CpG shores and CpG shelves, and a vast majority of CpG dinucleotides situated in the inter-CG island (inter-CGI) regions, also known as open sea (**Figure 2**). The mapping of long reads carried out with minimap2 or ngmlr did not affect the distribution of CpG sites according to the respective genomic annotations. Contrary to the results produced by nanopore sequencing, the 450K microarray and DREAM methodologies showed a tendency for CpG annotations in CpG islands, shores and shelves. This is due to the fact that, in the former, the CpG sites were chosen manually when the array probes were designed and, in the latter, there is a bias towards used enzyme restriction sites in those regions.

**Figure 2.**
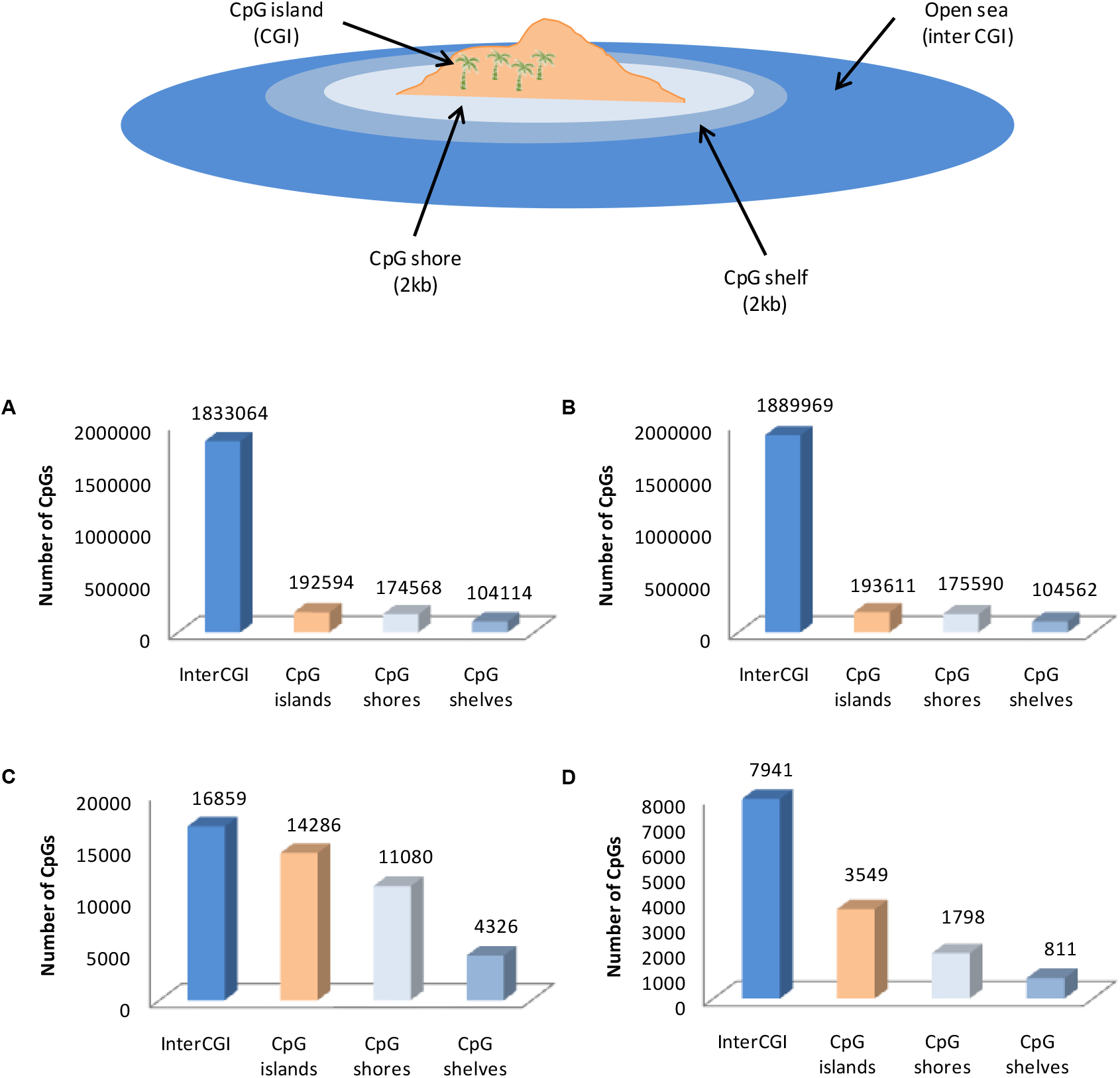
Number of chromosome 1 CpG sites interrogated in the different methodologies according to their genomic annotation (CpG islands, shores and shelves, and open sea regions) in the hg38 human genome reference sequence. **A**, nanopore_ngmlr; **B**, nanopore_minimap2; **C**, 450k microarray; **D**, DREAM. Whole genome nanopore sequencing produced a more consistent distribution of CpG sites in accordance with the expected sizes of the annotated regions, whereas 450k microarray and DREAM showed a more biased distribution of CpGs towards CpG islands, shores and shelves.

### Correlation analysis of the methylation status of CpG sites between nanopore sequencing, 450k microarray and DREAM

We performed a correlation analysis between the methylation frequencies obtained in this study with the corresponding values obtained in the 450k microarray and DREAM studies for the HEL cell line. The workflow that was used to compare the various data sets is represented as a diagram in **Figure 3**. The methylation status of each CpG site is calculated for each method using different calculation formulas and can vary between 0 (each copy of the CpG site is unmethylated) and 1 (each copy of the CpG site is methylated). In nanopolish, the frequency of methylation is calculated for each CpG site as “called_sites methylated”/“total called sites”. In the case of DREAM, we used an equivalent approach, that is, we calculated the frequency of methylation as the “number of methylated reads”/“number of total reads” that cover each CpG site. In the case of 450k microarray, the beta-value was used to quantitatively describe the methylation status of each CpG site, using the following formula: Beta-value = “methylated_signal intensity” / (“methylated_signal_intensity” + “unmethylated_signal_intensity” + α), where α (which has a fixed value of 100) is used when the intensities of the methylation and non-methylation signals are low^26^. Although the beta-value does not constitute a methylation frequency, as well as the M-value which is also used for calculating the methylation of CpG sites in arrays^42^, it can be interpreted as the proportion of methylated fragments in a given CpG site and, in this way, be legitimately compared with the methylation frequencies calculated based on the sequencing data.

**Figure 3.**
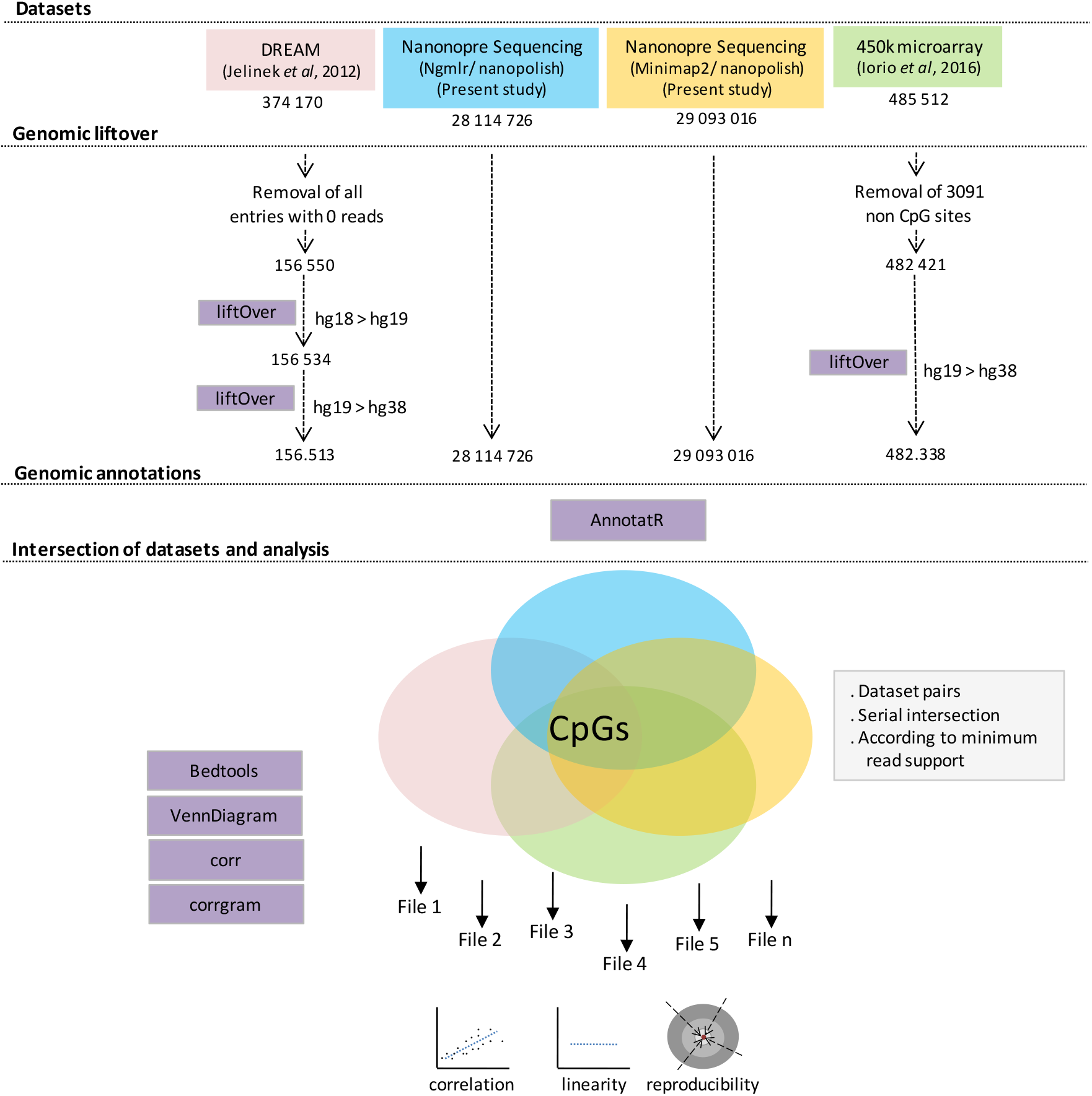
Diagrammatic representation of the general workflow used for benchmarking 5’-mC methylation values obtained by nanopore sequencing versus 450k microarray and DREAM. The numbers indicate the total number of CpG sites available at each step within the workflow. The analysis tools (liftOver, Bedtools and R) used in the workflow are represented by purple boxes.

Initially, the methylation frequencies produced based on the mapping of nanopore reads, using minimap2 or ngmlr, and the methylation values obtained by the 450k microarray or DREAM, were correlated with each other at the level of whole genome. For a total of 28,098,525 common CpG dinucleotides mapped by minimap2 and ngmlr (which include 99.9% of CpGs mapped by ngmlr and 96.6% of CpGs mapped by minimap2), the Pearson correlation was 0.98 (**Table II**), showing that the two mappings do not globally impact the results of methylation-calling carried out by nanopolish. The intersection of the methylation frequency files obtained based on ngmlr and minimap2, with the 450k microarray data file, produced a total of 478,633 and 480,281 common CpGs, corresponding to 99.2% and 99.6% of the CpGs contained in the methylation array, respectively. In the case of DREAM, after filtering the sites that did not have methylation values, and converting the genomic coordinates from hg18 to hg19, and then from hg19 to hg38, the intersection with the data obtained by nanopore sequencing, based on ngmlr and minimap2, resulted in a total of 148,346 and 152,686 common CpGs, corresponding to 94.8% and 97.6% of the CpGs contained in DREAM, respectively. The lower percentage of overlap in the coordinates of DREAM in relation to those of the methylation array, may be due to the fact that those correspond to a very old version of the human genome reference sequence (hg18). This implied that two rounds of conversion between genome coordinates have been performed in DREAM instead of 1 in the case of methylation array, which potentially led to a lower number of correctly converted CpG sites in the former. In both cases, very high correlations were observed between the methylation values of nanopore sequencing and 450k microarray (0.85), and between the methylation values of nanopore sequencing and DREAM (0.80).

**Table II.** Pearson correlations between nanopore sequencing, 450k microarray and DREAM (grey cells) and corresponding number of interrogated CpG sites (white cells). The p-value obtained in each correlation was < 2.2e-16.

|  | nanopore_ngmlr | nanopore_minimap 2 | 450k microarray | DREAM |
| --- | --- | --- | --- | --- |
| nanopore_ngmlr | 28 114 726 | 0.98 | 0.84 | 0.79 |
| nanopore_minimap 2 | 28 098 525 | 29 093 016 | 0.85 | 0.80 |
| 450k microarray | 478 633 | 480 281 | 482 338 | 0.81 |
| DREAM | 148 356 | 152 686 | 5 714 | 156 513 |

These correlations could suggest that these were the result of comparing, on one hand, a different number of CpGs in each case and, on the other hand, a significant proportion of different CpG sites, as a consequence of the intersection between the different datasets. To test this possibility, the coordinates of the CpG sites of the various datasets were serially intersected until a set of 5416 common CpG sites was obtained. The resulting correlations calculated for each paired datasets revealed that most values were similar to those obtained previously, indicating that the methylation values are well correlated even when comparing only ~0.02% of the total CpGs in the human genome (**Figure 4**). Likewise, the nanopore data obtained with the minimap2-mapped reads showed correlation values slightly higher than those obtained with ngmlr. In contrast, DREAM produced slightly stronger correlation values than the 450k microarray when compared to nanopore data. One possible explanation for this finding is a sample effect size, since the 5416 CpGs represent 3,5% of CpGs present in the DREAM data in contrast to only 1,1% of those present in the microarray. Taken together, these data indicate that methylation-calling based on low coverage nanopore sequencing produces results comparable to those obtained with other methodologies.

**Figure 4.**
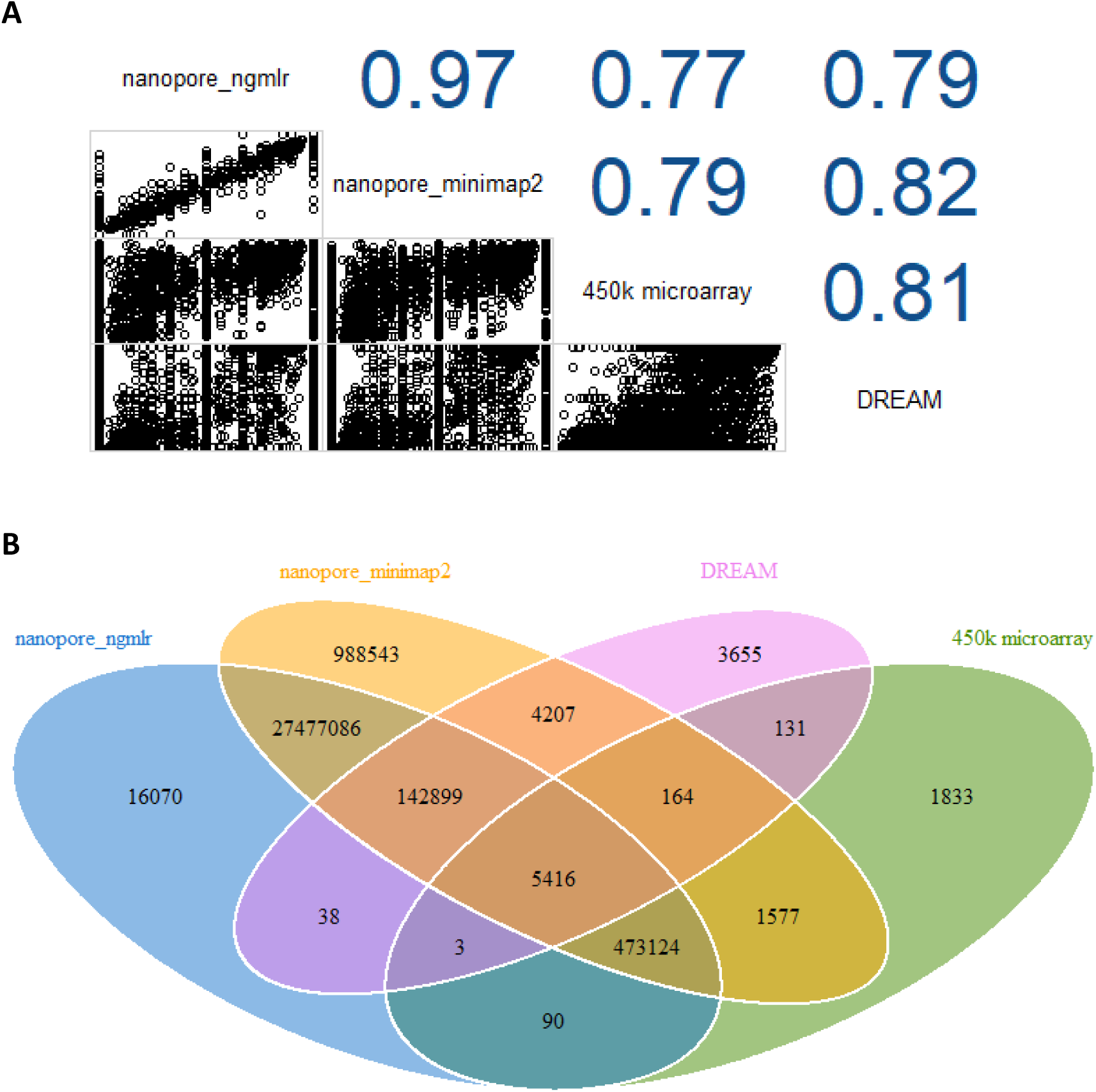
Correlation analysis of methylation values between each pair of methodologies. The CpG sites interrogated by nanopore_ngmlr, nanopore_minimap2, 450k microarray and DREAM, were intersected in a serial manner to generate a set of 5416 CpG sites common to all datasets. **A**, Correlation diagram indicating the Pearson correlation values and the corresponding plots for each pair of methodologies. **B**, Venn diagram showing the number of CpG sites resulting from the intersection of the various datasets. The plots were generated using packages corrgram (v1.13) and VennDiagram^51^ (v1.6.20) in R.

### Correlation of methylation values as a function of read support and depth

Then, we asked if the correlation of methylation values between nanopore sequencing and the other methods could be stronger as a result of an increase in minimum coverage depth. To simulate this effect, we filtered the positions of CpG sites in the methylation frequency files according to a read support between 2X and 20X, at 1X unit intervals. Pearson correlation was calculated for each minimum coverage level and plotted as a function of read support (**Figure 5**). This analysis showed that the correlation between nanopore and 450k microarray data increased up to 0.89 and 0.90 at 17X read support, using minimap2- and ngmlr-based methylation frequencies, respectively. From this point on, the corresponding correlation values began to decrease until the maximum read support of 20X. This likely results from the fact that the number of available CpGs approaches zero when the read support is highest and that the corresponding methylation frequencies are calculated on likely incorrectly mapped reads or secondary read alignments.

**Figure 5.**
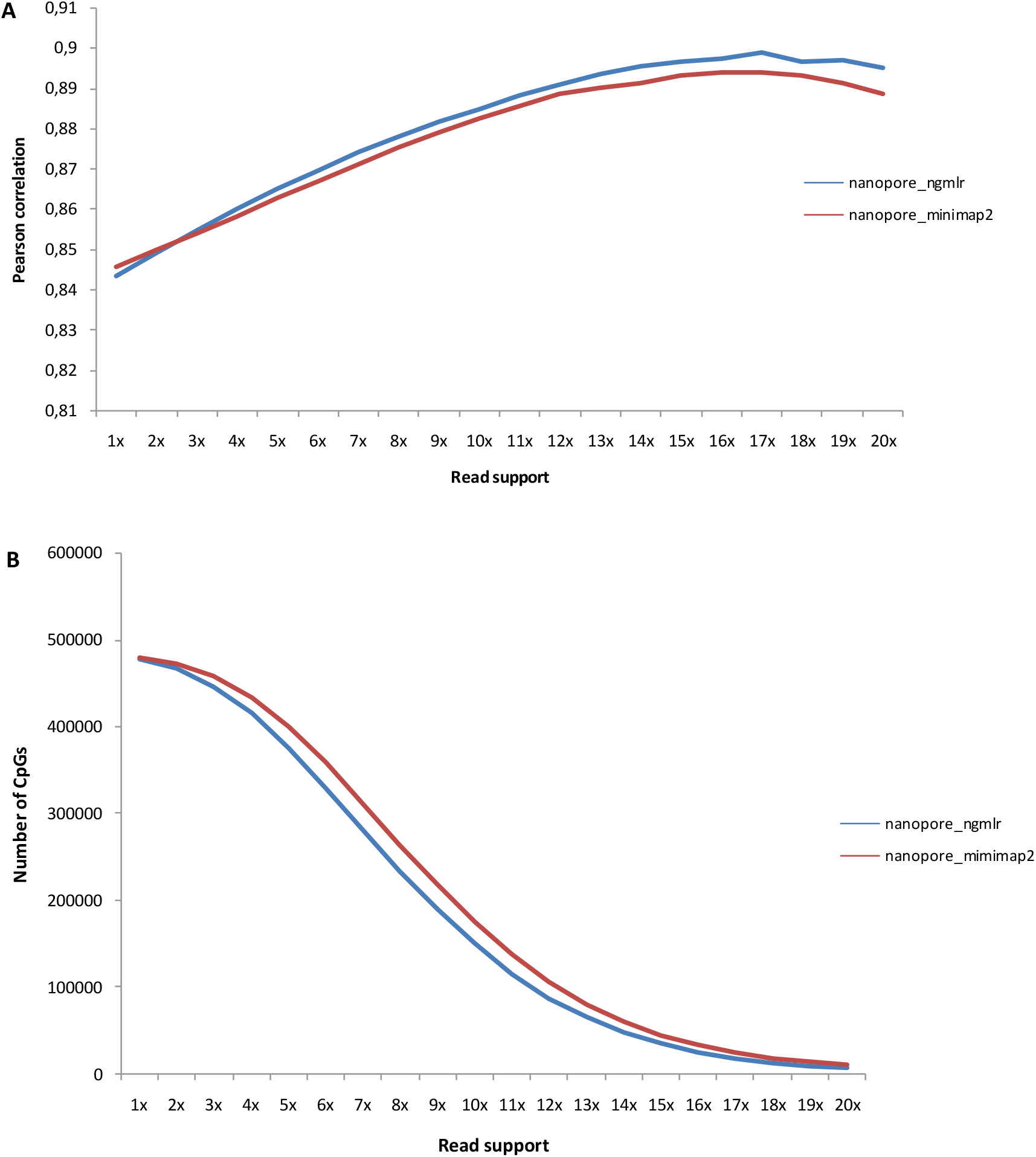
Evolution of Pearson correlation values as a function of minimum read support. The CpG sites mapped with minimap2 and ngmlr were filtered from the methylation frequencies file according to a minimum read support between 1X and 20X at a 1X intervals. **A**, Pearson correlation values as a function of read support. **B**, Number of CpG sites as a function of read support. The Pearson correlation values was based on the intersection of genome-wide CpG sites between each of the nanopore datasets and the 450k microarray dataset. The correlation increased up to 17X read support. From 18X to 20X the correlation values showed a slight decrease likely as a consequence of a very small number of interrogated CpG sites.

Conversely, we also sought to see the impact of a diminished average depth of coverage on the correlation values. For this purpose, we analyzed separately the data obtained in each of the 4 sequencing runs, which produced an average coverage depth of 1.8X/1.7X, 2.3X/2.2X, 2.5X/2.4X and 4.0X/3.8X, using minimap2/ngmlr, respectively. The total number of CpGs mapped is directly proportional to the average depth of coverage and the size of the intersection CpG sets is also proportional to the correlation values between each pair of runs. (**Table III**). When the number of CpGs was lowest, the correlation between nanopore data (based on minimap2-mapping) and DREAM or 450k microarray was of 0.73 or 0.76, respectively (**Table IV**). The correlation increased as a function of the number of available CpGs, reaching the maximum values of 0.76 and 0.79, respectively. These results indicate that an average coverage depth of 1-2X in nanopore sequencing can provide reliable information on the methylation status of a majority of CpG dinucleotides.

**Table III.** Pearson correlations between each 2 runs of nanopore sequencing (grey cells) and corresponding number of interrogated CpG sites (white cells). The CpGs were called based on minimap2 mapping results. The p-value obtained in each correlation was < 2.2e-16.

|  | Run 1 | Run 2 | Run 3 | Run 4 |
| --- | --- | --- | --- | --- |
| Run 1 | 20 517 732 | 0.69 | 0.70 | 0.72 |
| Run 2 | 16 430 158 | 22 667 288 | 0.71 | 0.73 |
| Run 3 | 17 166 563 | 18 887 356 | 23 751 970 | 0.74 |
| Run 4 | 19 126 896 | 21 077 222 | 22 071 529 | 26 794 587 |

**Table IV.** Pearson correlations between each run of nanopore sequencing and 450k microarray or DREAM, and corresponding number of interrogated CpG sites. The CpGs called in nanopore sequencingnwere based on minimap2 mapping data. The p-value obtained in each correlation was < 2.2e-16.

|  | 450k microarray |  | DREAM |  |
| --- | --- | --- | --- | --- |
|  | CpGs | Pearson correlation | CpGs | Pearson correlation |
| Run 1 | 346 550 | 0.76 | 93 294 | 0.73 |
| Run 2 | 382 218 | 0.76 | 104 657 | 0.74 |
| Run 3 | 408 422 | 0.78 | 115 047 | 0.74 |
| Run 4 | 447 650 | 0.79 | 130 951 | 0.76 |

### Reproducibility and linearity of methylation frequency data

When implementing nanopore sequencing technology to study 5’-mC, it is important to evaluate the reproducibility of the data. For this purpose, we consecutively intersected the coordinates of the CpG sites aligned with minimap2 in the 4 sequencing runs, and correlated the corresponding methylation frequencies between each pair of runs, so that the various correlations were directly comparable. Moreover, in order to avoid comparing CpG sites with a very different number of mapped reads, due to the different average coverage depth obtained in each run, we used only the methylation frequencies of the sites that were supported by a minimum of 5 reads, resulting in a total of 255 176 CpG sites. Although the average coverage depth in the 4 runs was very low (1.8-4.0X), strong correlation values were still obtained between different sequencing runs (**Figure 6**).

**Figure 6.**
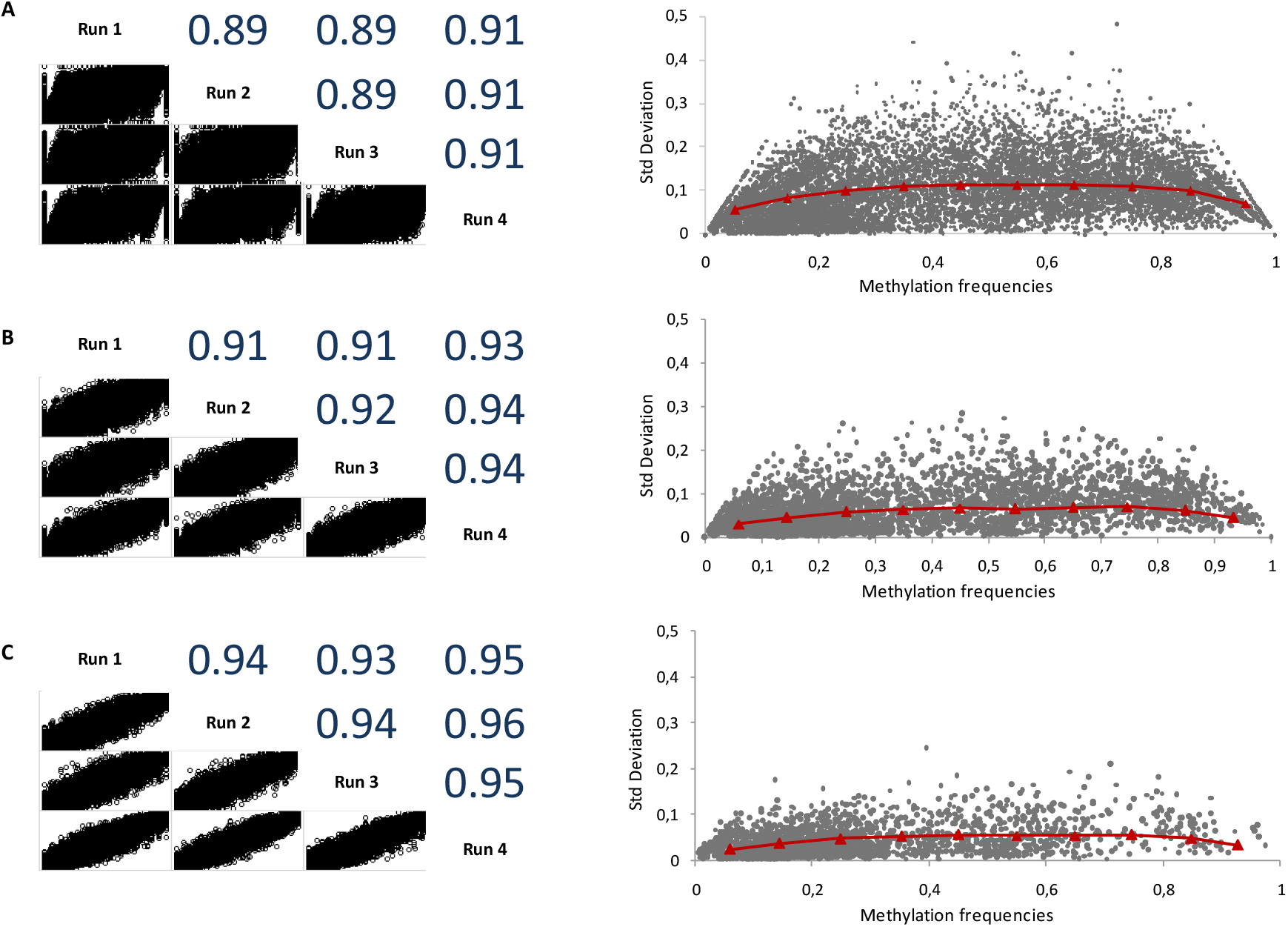
Representation of the correlation values and distribution of methylation frequency variation obtained in technical replicates, as a function of minimum read support. **A**, 5X read support; **B**, 10X read support; **C**, 15X read support. The median of the standard deviations was calculated for each of 10 bins between 0 and 1, and is represented by small red triangles connected by red lines in each of the plots on the right side of the figure. The linearity of the variation in methylation frequencies as well as the correlation values increase as a function of read support.

The existence of technical replicates of the sequencing runs also allowed us to analyze the linearity of the methylation frequency data between 0 and 1. The purpose of this analysis was to determine if the low depth of coverage introduced a bias in the methylation frequencies in the ends of the scale. For this aim, we used the sets of 255 176, 81 287 and 53 083 CpG sites common to all runs and that were supported by at least 5, 10 or 15 reads, respectively. For each of these sites, we calculated the mean and standard deviation of the methylation frequencies. Then, we calculated the median of the standard deviations for each of 10 bins between the values of 0 and 1 and plotted these values against the scatter plot of mean versus standard deviation of the methylation frequencies. The results showed that there is an almost linearity of the data between 0 and 1 with only a slight compression at the ends. The difference between the maximum and minimum median values at 15X was half than that at 5X read support (0,03 versus 0,06 respectively), indicating that complete linearity may be reached at high depths of coverage.

### Distribution of methylation frequencies in distinct genomic contexts

We then carried out a finer analysis of the methylation values according to the genomic context of the CpG sites. This analysis allows a biological insight into the data produced by the different methodologies and could help to identify specific differences in the methylation profiles between the various datasets. For this end, we compared the distribution of methylation values in each methodology using 21 bins between 0 and 1. We chose to use only the CpGs on chromosome 1 as representative of the distribution of methylation values at the genome level, in order to reduce the computational effort associated with analysis of the complete datasets. Comparative analysis of the methylation frequency profiles obtained with minimap2- and ngmlr-based nanopore sequencing data showed an almost complete overlap, reflecting the very strong correlation of methylation frequencies (**Figure 7**). The DREAM profile, which is also based on sequencing data, revealed a similar global profile to nanopore data. In contrast, the 450k microarray profile showed a more balanced distribution of methylation values between 0 and 1, with a decrease in both extreme bins compared to the profiles obtained with both sequencing approaches.

**Figure 7.**
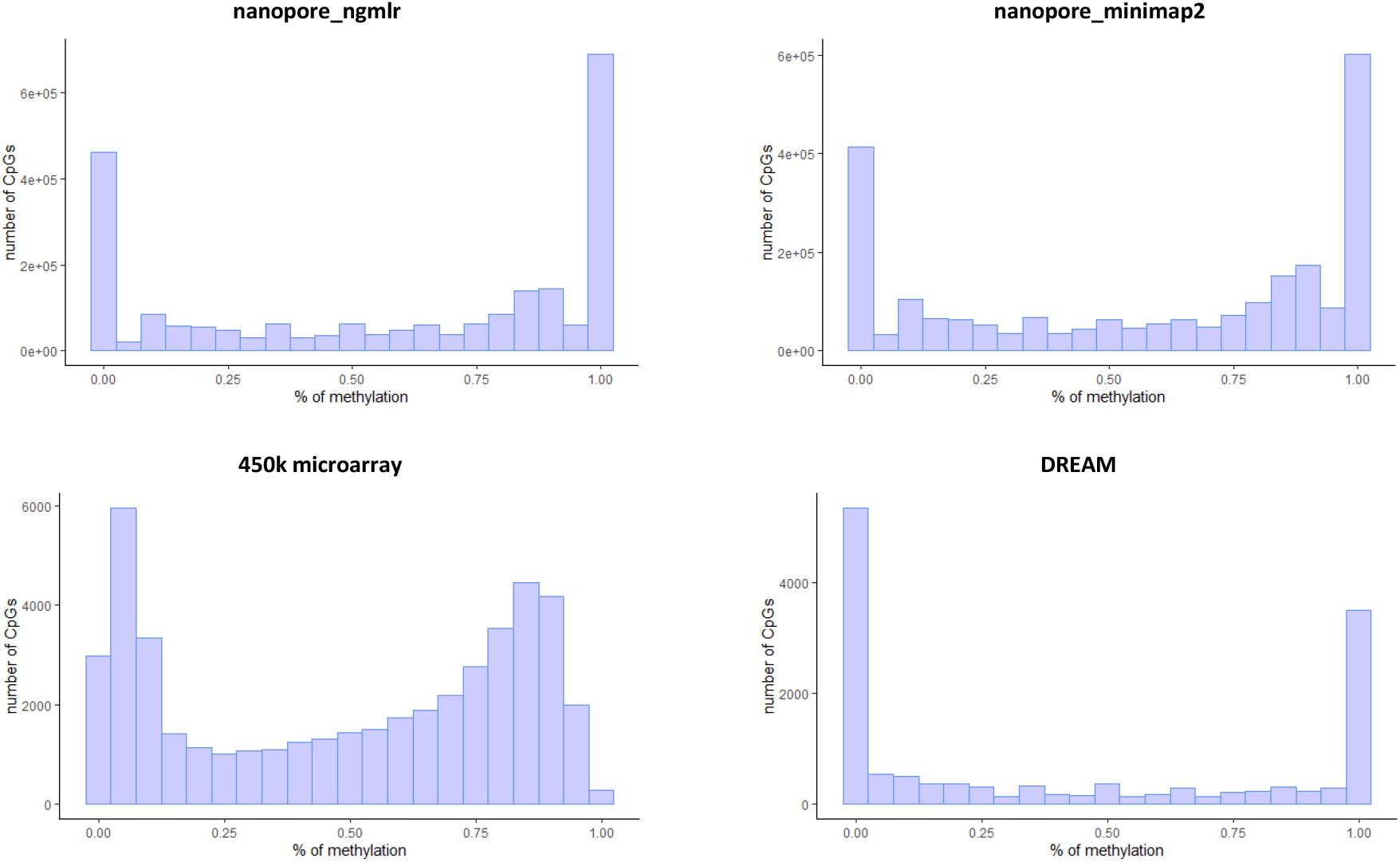
Distribution of methylation values for chromosome 1 CpGs obtained in nanopore sequencing, 450k microarray and DREAM. Plots were generated using ggplot2^52^ (v3.3.2) R package. Both sequencing approaches show a predominance of fully unmethylated and full methylated CpG sites compared to the 450k microarray where the distribution of methylation frequencies is more balanced along the scale.

We subsequently broke down the methylation values according to the genomic context in order to detail the differences observed in the global genomic profiles. We observed in the various methodologies a continuum in the variation of the distribution of methylation values from CpG islands to inter CGI regions. In CpG islands, the vast majority of CpGs are in an unmethylated state whereas in the CpG shelves and inter CGI regions there is a predominance of methylated CpGs (**Figure 8**). In contrast, there is a more balanced distribution between complete unmethylated and complete methylated sites in CpG shores among all methodologies. In particular, nanopore sequencing showed a tendency towards a predominance of methylated CpGs whereas DREAM revealed the opposite scenario. This profile analysis also emphasized some details about the calculation of methylation in the various methodologies. In the case of DREAM, almost 43% of all mapped CpG sites have 5 or less support reads (data not shown), indicating that for many CpGs there was not enough vertical coverage to sufficiently sample each site. The consequence of this is that there may be an excess of completely methylated and unmethylated sites, relative to sites where there is co-existence of methylated and unmethylated CpGs. In the 450k microarray, the number of CpG sites in the extreme bins is much lower than the corresponding bins in the other methodologies. This difference is due to the fact that the Infinium II HumanMethylation450 methylation array assays demonstrate a right shift for fully unmethylated CpGs and a left shift for fully methylated CpGs, that is, a compression of beta-values towards the center of the scale^26^. Taken together, low coverage depth nanopore sequencing provides a reliable global methylation profile of the human genome.

**Figure 8.**
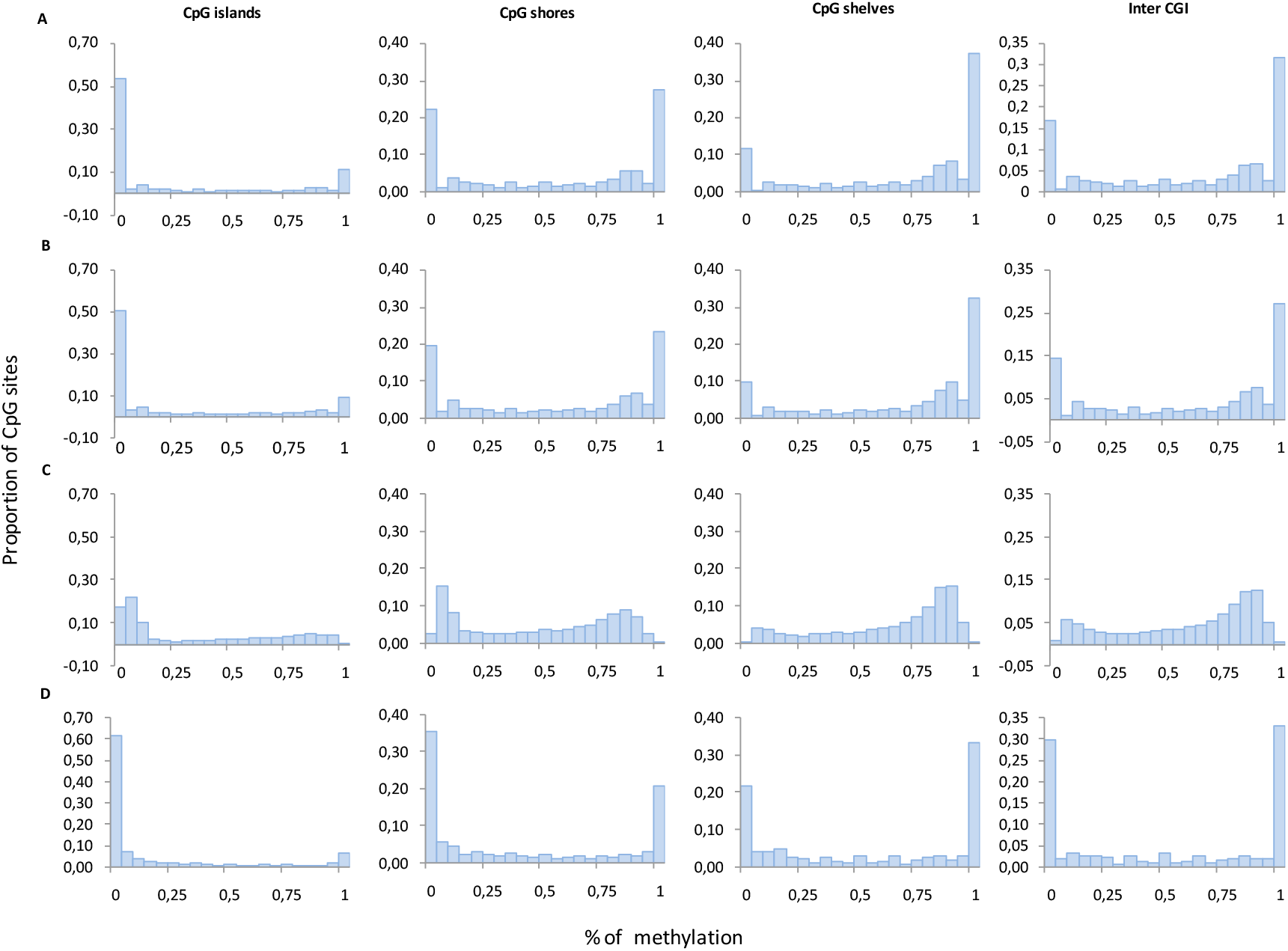
Distribution of methylation values obtained in nanopore sequencing, 450k microarray and DREAM, according to the CpG genomic annotations. **A**, nanopore_ngmlr; **B**, nanopore_minimap2; **C**, 450k microarray; **D**, DREAM. In CpG islands the majority of interrogated CpGs are unmethylated. In CpG shores, there is a balanced distribution between unmethylated and methylated CpG sites, whereas in CpG shelves and inter-CGI regions there is a predominance of methylated CpG sites.

## Discussion

Nanopore sequencing is a technology that emerged after the widespread use of microarray and short-read sequencing technologies in the scientific community. For this reason, several sequencing applications, including the study of 5’-mC, were already well established when it was shown, for the first time, that nanopore sequencing was able to detect 5’-mC in the human genome using the MinION device. Despite the fact that this device is not yet prepared to generate a large amount of sequencing data, it is nonetheless worthwhile exploring the capabilities of MinION to generate whole genome 5’-mC data from sequencing at low coverage depth. In this context, we proposed to carry out a benchmark study to compare the performance of nanopore sequencing with methylation microarray and DREAM in the study of methylation in the human cell line HEL. To do this, we performed whole genome sequencing in the MinION, mapped the reads in the human reference sequence using 2 different mapping tools, performed 5’-mC methylation-calling using nanopolish, and compared the methylation values between the different methodologies using various statistical tools. The sequencing data produced by MinION showed a significant proportion of long high quality reads, which allowed us to assess the methylation level of almost all CpG sites existing in the human genome sequence.

At a depth of coverage of approximately 10X, the correlation between the methylation frequencies obtained by nanopore sequencing and the methylation values obtained by DREAM or the 450k microarray, was approximately 0.80 and 0.85, respectively, regardless of the tool used in the mapping of long reads. The simulation of higher coverage depths, based solely on CpG sites with a minimum number of support reads, confirmed the progressive and linear increase of correlation up to 0.90, demonstrating that methylation-calling based on low coverage depth nanopore sequencing provides reliable methylation frequency data.

Conversely, we showed that even at a coverage level <2X, the lowest correlation obtained was 0.73 between nanopore sequencing and DREAM. Although strong correlations have been obtained between the various methylation data sets in all cases, it is important to emphasize that exposure to different cell culture conditions, as well as the use of different methods for the extraction and preparation of genomic DNA, carried out in distinct laboratories at different times, may have contributed to small changes in the methylation pattern of the HEL cells.

Using 4 different library preparations and nanopore sequencing runs, we observed a very strong correlation between methylation frequencies of the HEL cell line obtained in any of 2 assays. In addition, we observed an almost linearity of the methylation frequencies between the minimum (fully unmethylated CpGs) and maximum (fully methylated CpGs) values at low levels of read 0-0.2 support. These results contrast with the intrinsic lack of homoscedasticity reported to occur in the and 0.8-1.0 intervals of beta-values in methylation arrays^42^.

The methodologies for studying 5’-mC in the human genome have different advantages and disadvantages. In the case of 450k microarray, genomic DNA is initially treated with bisulfite, which converts unmethylated cytosines to uracils. This converting action may not be completely efficient, leaving some unconverted cytosines in an unmethylated state which are later interpreted as being methylated. This phenomenon is particularly relevant when the genomic DNA is not fully denatured before bisulfite treatment, because only the cytosines that are found in single-stranded DNA are susceptible to conversion^15^. In addition, the high temperature (99°C) used in some bisulfite treatment protocols can lead to the degradation of genomic DNA, which is manifested by random breaks in the sequence as a result of depurinations^16^. These breaks can give rise to low molecular size fragments, which can be lost in the next steps of the procedure, reducing the sequence representativeness of the original sample. Before hybridization on the chip, additional steps of sample manipulation can also cause a representation bias through the phenomena of preferential sequence selection during PCR, loss of amplified fragments during purification and incomplete denaturation of the DNA. Moreover, bisulfite treatment cannot distinguish between 5’-mC and 5’-hydroxylmethylcytosine epigenetic modifications. Bisulfite transforms 5’-hydroxylmethylcytosine into cytosine-5-methylsulfonate, which is read as a C during sequencing, making the presence of cytosines indistinguishable with either form of modification^17^.

Bisulfite treatment is also used in short-read whole-genome bisulfite sequencing (WGBS)^43^, which can be affected by a similar set of conditions as the methylation array. Moreover, in short read sequencing technologies such as Illumina, sequence diversity is one of the critical aspects that allows high quality sequences to be obtained. In WGBS, unmethylated cytosines are converted into uracils, thus artificially reducing sample diversity and impacting overall sequence quality. Short reads, such as those used in the DREAM method, are also less adapted than long reads to map repetitive and highly homologous regions that exist in the human genome. Compared to the methylation array and DREAM, which only query a small fraction of CpG sites distributed essentially by gene regions and CpG islands, nanopore sequencing and WGBS offer a global view of the methylation profile of the human genome. The analysis of CpGs at a global level translates into an important advantage in terms of knowledge of epigenetic mechanisms, as it may allow the detection of CpG sites with novel functional relevance. In addition to the long read capabilities of nanopore sequencing, genomic DNA can be sequenced in its native form without any type of in vitro modification that can alter its integrity, sequence and representativeness, thus maintaining the original epigenetic landscape of the sample. However, to achieve the desired sequencing coverage that we obtained in this study, there may be necessary a few rounds of sequencing, thus impacting on the overall cost of nanopore sequencing compared to the other traditional methodologies. However, the higher cost may be partially overcome with future improvements on sample quality, library preparation and/or sequencing chemistry, thereby being able to generate a larger amount of sequence data with less sequencing runs.

Another important advantage of nanopore sequencing is the fact that it has the possibility of following novel scientific developments in the field of epigenomics, without necessarily implying upgrades in the technological platform or changes in the sequencing methods. This reflects the fact that the different types of epigenetic modification of DNA or RNA, including the various forms of methylation, can in theory be detected using the same primary data, without the need to repeat the sequencing of the samples. To achieve this, several computational methods have been developed capable of identifying different types of epigenetic modifications^44^. These can generally be divided into 2 main types, which include methods based on statistics and methods based on models. The former do not require the existence of a training model for DNA or RNA modifications, but they require the sequencing of a matched control sample to detect the various types of methylation^45,46^. Non-statistical methods can further be divided into mapping-dependent methods^21,47–49^ and basecalling-only methods such as Guppy^31^ or Flappie^50^ (https://github.com/nanoporetech/flappie). Model-based methods may be more advantageous as they reduce the extra costs of sequencing a control sample, but they need to train a model based on the species genome, type of epigenetic modification and flow cell version. In this work, nanopolish has shown a good performance in calling 5’-mC in the human genome using R9.4 MinION flow cells. It is nonetheless desirable to carry out benchmarking studies using other methylation callers, in order to identify the best tool for determining 5’-mC in human genome samples. These tools should also be tested in the context of a comparative analysis between samples not exposed and exposed to an environmental agent, in order to assess the differences in the global methylation pattern. In summary, we showed in this study that methylation calling, based on nanopore sequencing, is a fast, robust and sensitive approach for determining the status of 5’-mC in the human genome.

## Data availability

The datasets presented in this study are available in the European Nucleotide Archive (ENA) at EMBL-EBI under the accession number PRJEB45078 (https://www.ebi.ac.uk/ena/browser/view/PRJEB45078).

## Acknowledgements

This work is a result of the GenomePT project (POCI-01-0145-FEDER-022184), supported by COMPETE 2020 - Operational Programme for Competitiveness and Internationalisation (POCI), Lisboa Portugal Regional Operational Programme (Lisboa2020), Algarve Portugal Regional Operational Programme (CRESC Algarve2020), under the PORTUGAL 2020 Partnership Agreement, through the European Regional Development Fund (ERDF), and by Fundação para a Ciência e a Tecnologia (FCT). This work was also supported by Fundos FEDER through the Programa Operacional Factores de Competitividade - COMPETE and by Fundos Nacionais through the FCT within the scope of the project UID/BIM/00009/2019 (Centre for Toxicogenomics and Human Health-ToxOmics).

## Notes

### Competing Interest Statement

The authors have declared no competing interest.

